## Supplemental Figures and video link for "Mouse germline cysts contain a fusome that mediates oocyte development"

### Supplementary Figures

Supplementary Videos link :

https://drive.google.com/drive/folders/1XuOoesa_WKMJUaVTpOKGbWWhn_fJc56M

### Fig. S1.

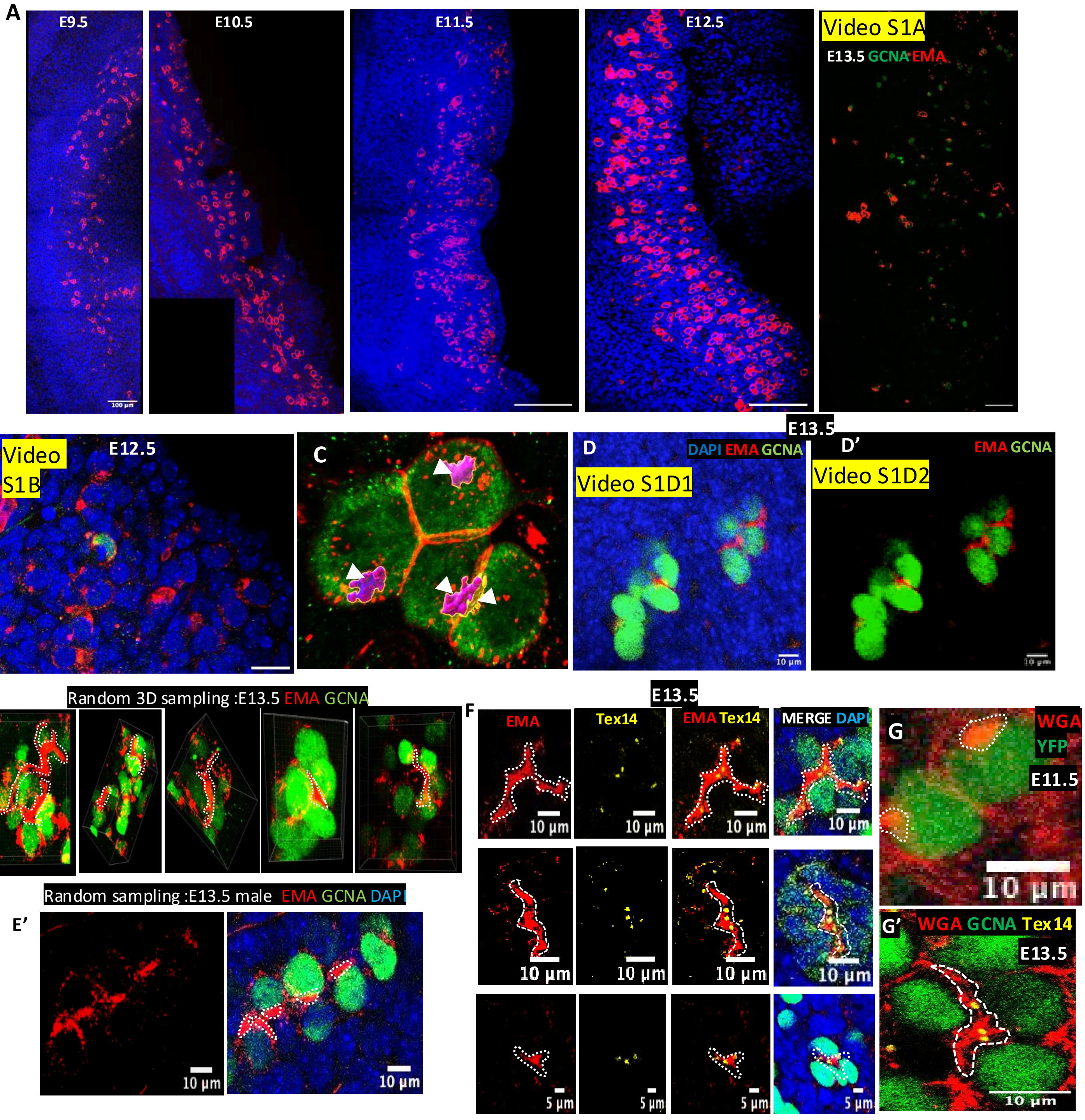

A
E9.5
100 µm
E10.5
E11.5
E12.5
Video S1A
E13.5 GCNA EMA
B
Video S1B
E12.5
C
D
DAPI EMA GCNA
Video S1D1
10 µm
E13.5
D'
EMA GCNA
Video S1D2
E
Random 3D sampling :E13.5 EMA GCNA
E'
Random sampling :E13.5 male EMA GCNA DAPI
10 µm
10 µm
E13.5
F
EMA
Tex14
EMA Tex14
MERGE DAPI
10 µm
5 µm
G
WGA
YFP
E11.5
10 µm
G'
WGA GCNA Tex14
E13.5
10 µm

Fig. S1 continued

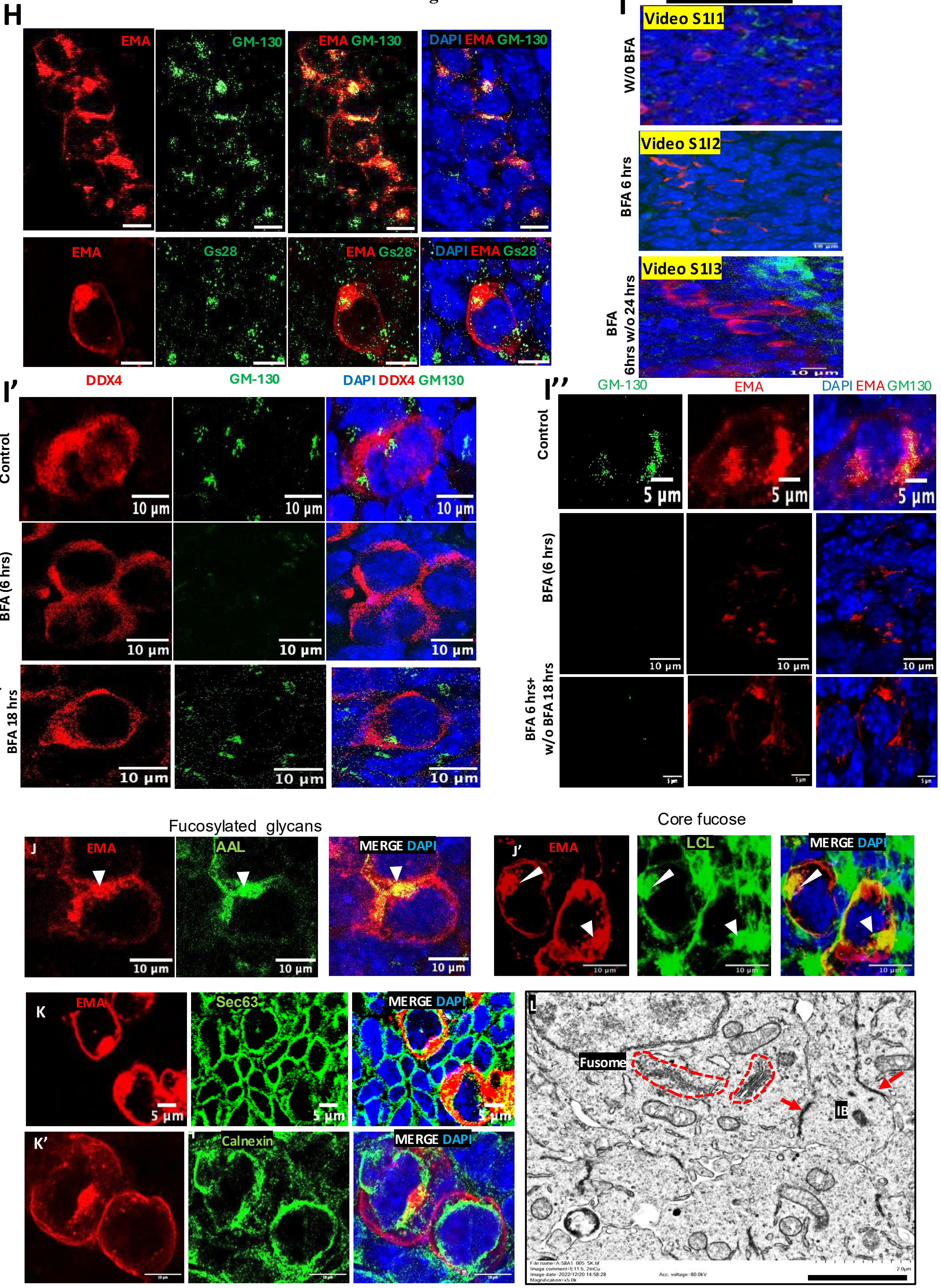

Figure S2

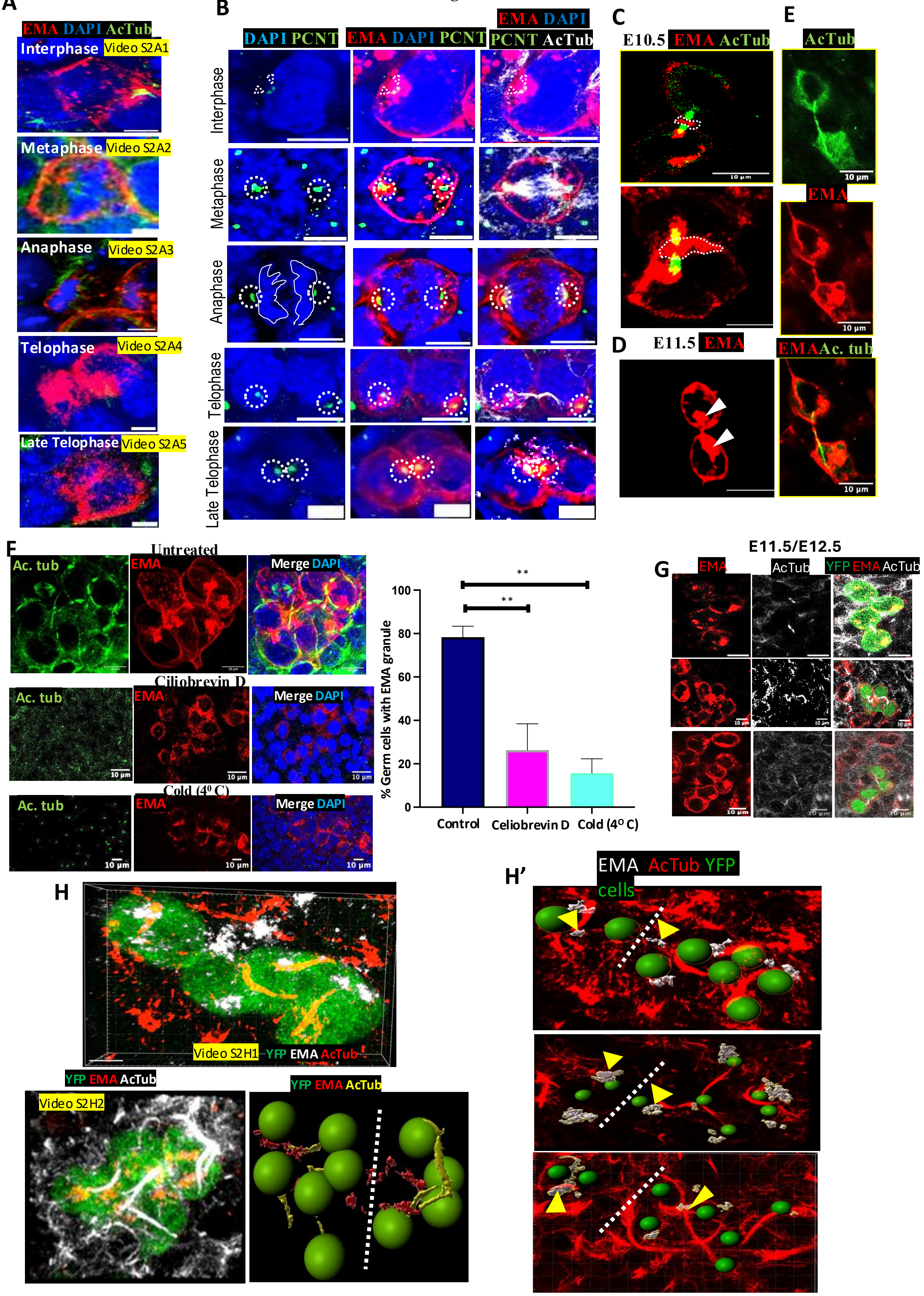
A
EMA DAPI AcTub
Interphase
Video S2A1
Metaphase
Video S2A2
Anaphase
Video S2A3
Telophase
Video S2A4
Late Telophase
Video S2A5
B
DAPI PCNT
EMA DAPI PCNT
EMA DAPI PCNT AcTub
Interphase
Metaphase
Anaphase
Telophase
Late Telophase
C
E10.5 EMA AcTub
10 µm
D
E11.5 EMA
E
AcTub
10 µm
EMA
10 µm
EMAAc. tub
10 µm
F
Untreated
Ac. tub
EMA
Merge DAPI
Ciliobrevin D
Cold (4⁰ C)
10 µm
100
80
60
40
20
0
% Germ cells with EMA granule
**
**
Control
Celiobrevin D
Cold (4⁰ C)
E11.5/E12.5
G
EMA
AcTub
YFP EMA AcTub
10 µm
H
Video S2H1
YFP EMA AcTub
YFP EMA AcTub
Video S2H2
YFP EMA AcTub
H'
EMA AcTub YFP cells

**Fig. S3.**

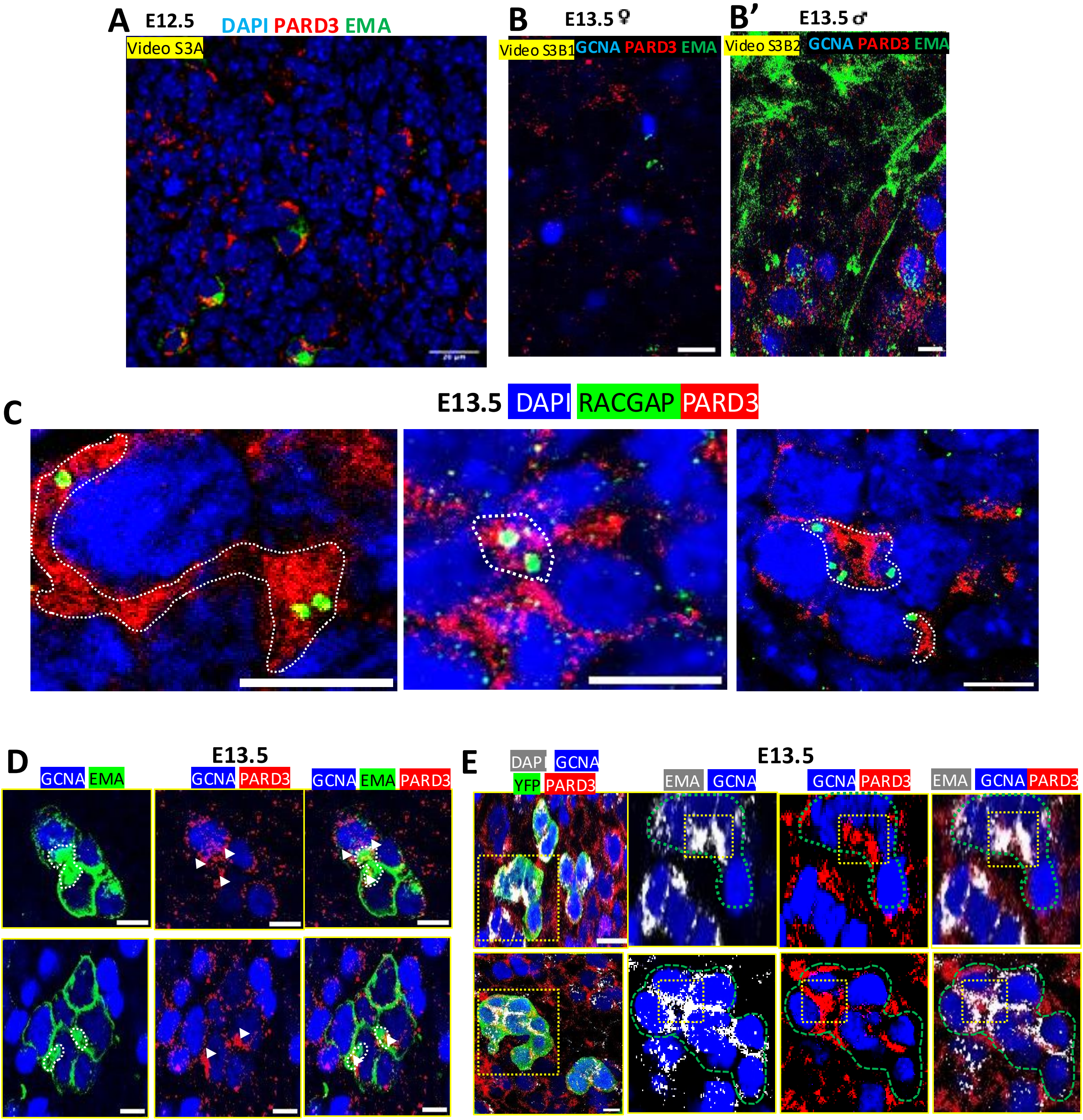
A
E12.5
DAPI PARD3 EMA
Video S3A
B
E13.5♀
Video S3B1
GCNA PARD3 EMA
B'
E13.5♂
Video S3B2
GCNA PARD3 EMA
C
E13.5 DAPI RACGAP PARD3
D
E13.5
GCNA EMA
GCNA PARD3
GCNA EMA PARD3
E
E13.5
DAPI GCNA YFP PARD3
EMA GCNA
GCNA PARD3
EMA GCNA PARD3

### Fig. S3 continued

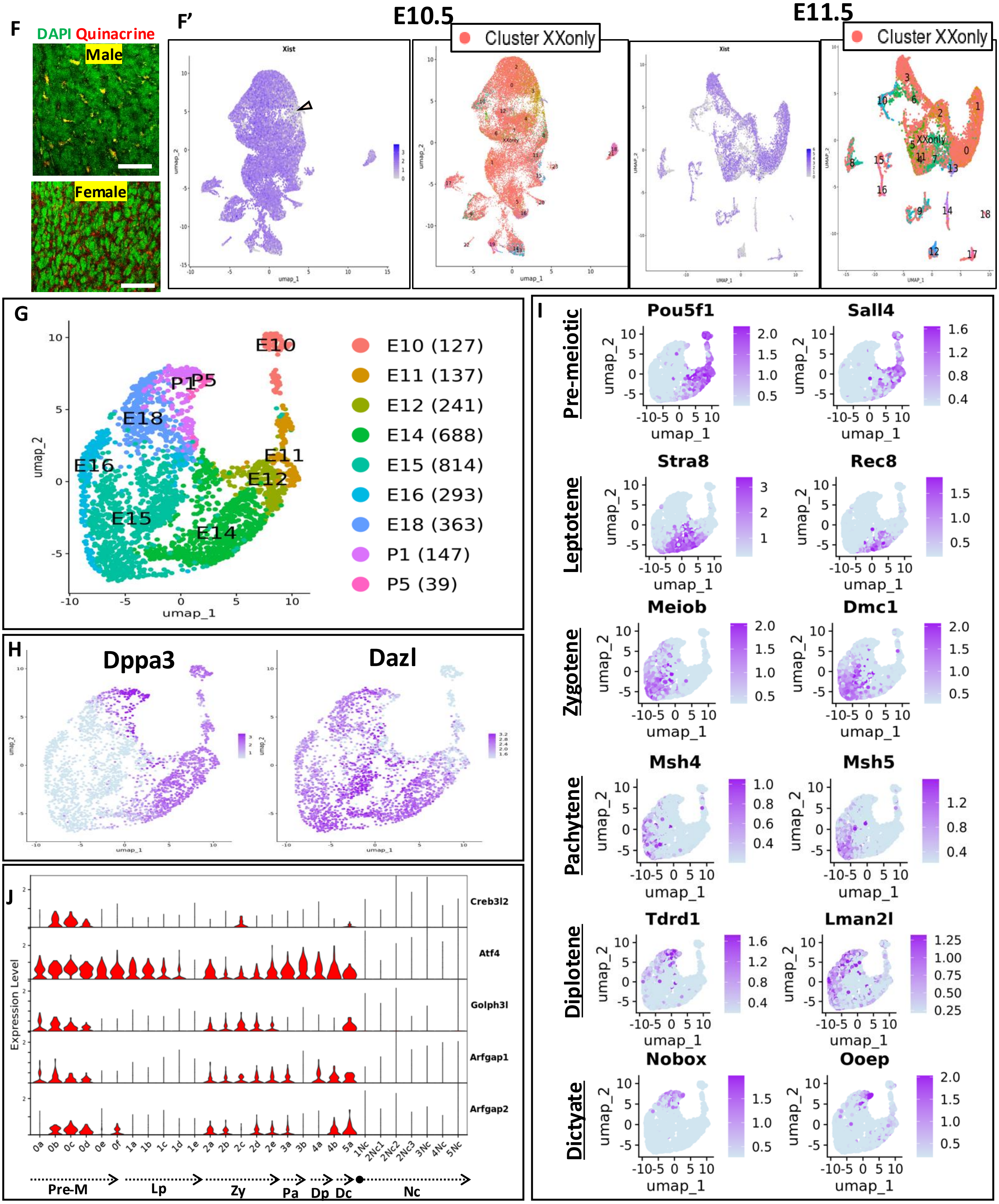

### Fig. S4

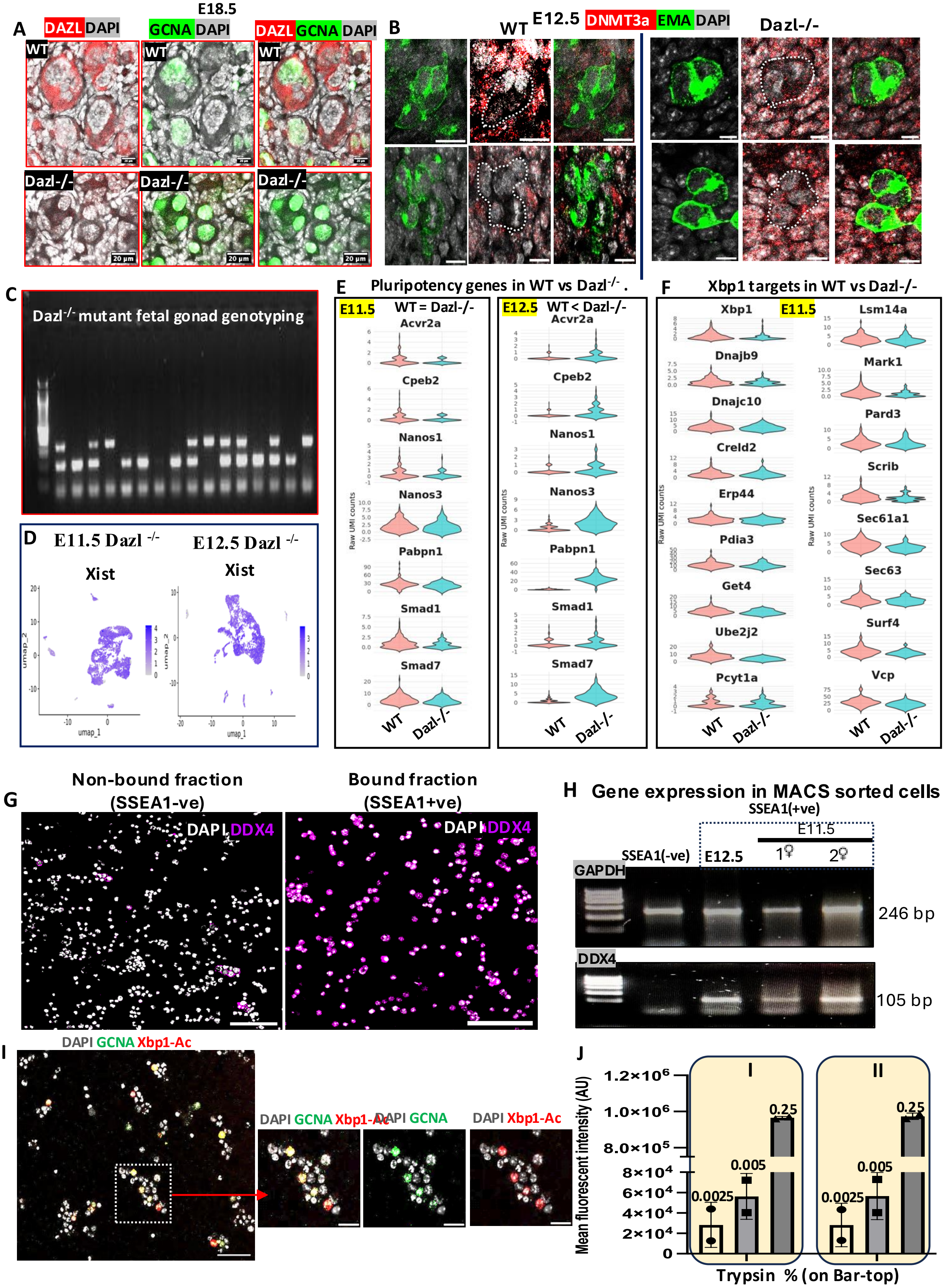
A
E18.5
DAZL DAPI
GCNA DAPI
DAZL GCNA DAPI
WT
Dazl-/-
20 µm
B
WT E12.5 DNMT3a EMA DAPI
Dazl-/-
C
Dazl-/- mutant fetal gonad genotyping
D
E11.5 Dazl -/-
E12.5 Dazl -/-
Xist
umap_1
umap_2
E
Pluripotency genes in WT vs Dazl-/- .
E11.5
WT = Dazl-/-
E12.5
WT < Dazl-/-
Acvr2a
Cpeb2
Nanos1
Nanos3
Pabpn1
Smad1
Smad7
Raw UMI counts
WT
Dazl-/-
F
Xbp1 targets in WT vs Dazl-/-
E11.5
Xbp1
Dnajb9
Dnajc10
Creld2
Erp44
Pdia3
Get4
Ube2j2
Pcyt1a
Lsm14a
Mark1
Pard3
Scrib
Sec61a1
Sec63
Surf4
Vcp
Raw UMI counts
WT
Dazl-/-
G
Non-bound fraction
(SSEA1-ve)
Bound fraction
(SSEA1+ve)
DAPI DDX4
H
Gene expression in MACS sorted cells
SSEA1(+ve)
E11.5
SSEA1(-ve)
E12.5
1♀
2♀
GAPDH
246 bp
DDX4
105 bp
I
DAPI GCNA Xbp1-Ac
DAPI GCNA
DAPI Xbp1-Ac
J
I
II
1.2×10⁶
1.0×10⁶
8.0×10⁵
8×10⁴
6×10⁴
4×10⁴
2×10⁴
0
0.25
0.005
0.0025
Mean fluorescent intensity (AU)
Trypsin % (on Bar-top)

### Fig. S5

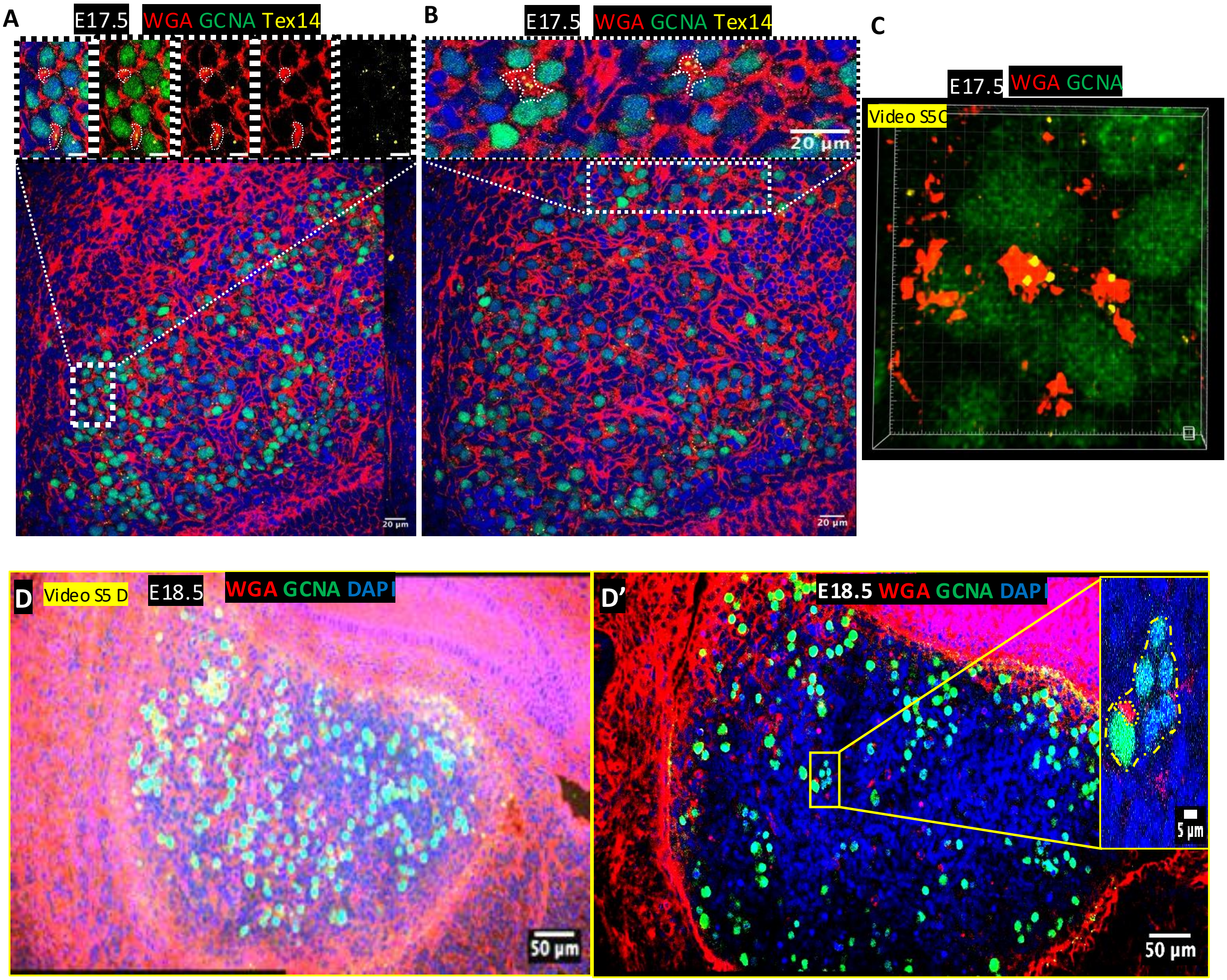
A
E17.5
WGA GCNA Tex14
20 µm
B
E17.5
WGA GCNA Tex14
20 µm
20 µm
C
E17.5
WGA GCNA
Video S5C
D
Video S5 D
E18.5
WGA GCNA DAPI
50 µm
D'
E18.5 WGA GCNA DAPI
5 µm
50 µm

### Fig. S5 continued..

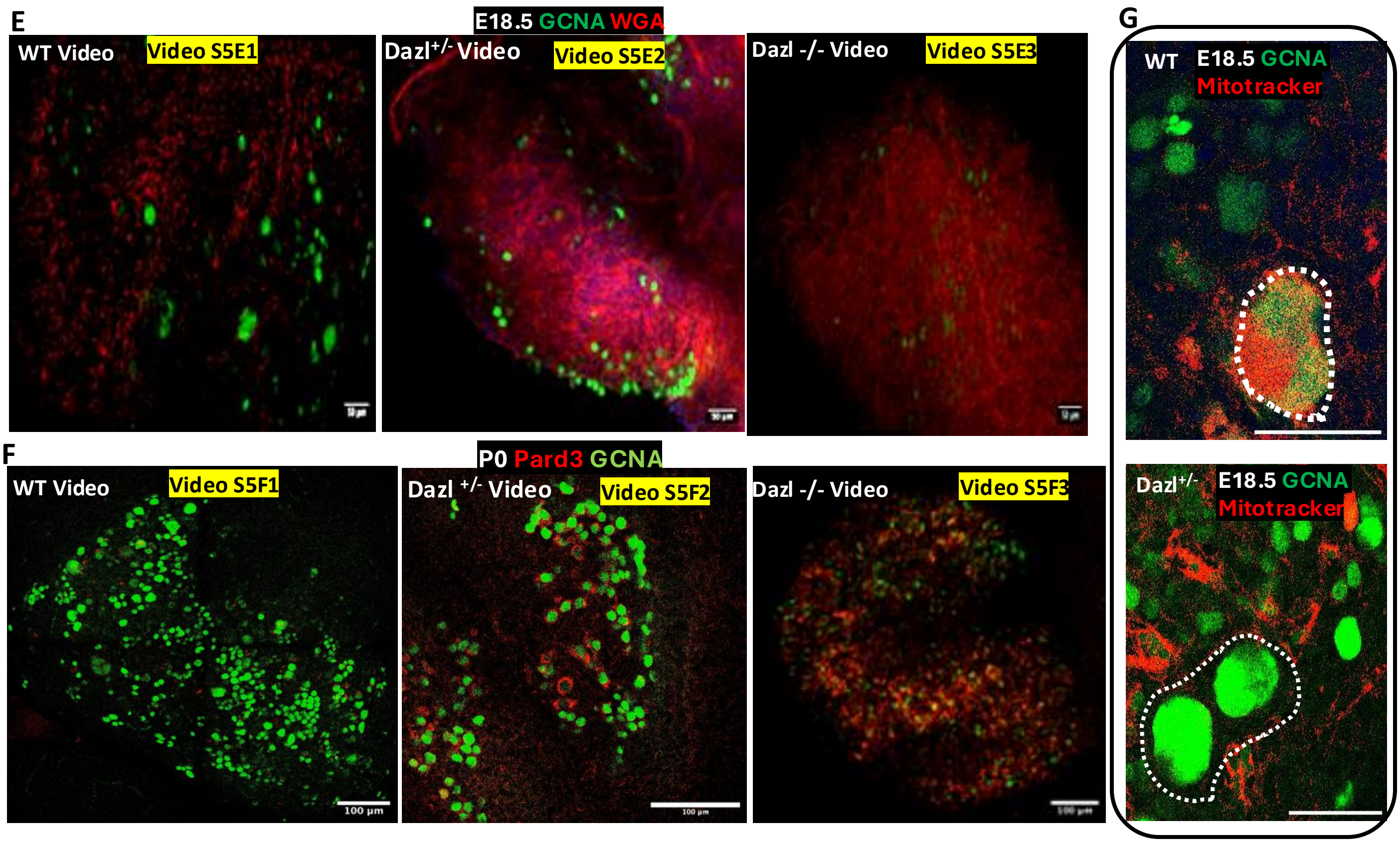
